# Compartment-Specific Assembly and Functional Potential of the Bacteriome of *Citrullus colocynthis* in a Semi-Arid Ecosystem

**DOI:** 10.64898/2026.06.05.730417

**Authors:** Khadija Ait Si Mhand, Salma Mouhib, Nabil Radouane, Imad khatour, Fatima-Zahra Aliyat, Mohamed Hijri

**Author notes:** Correspondence to Mohamed Hijri.

## Abstract

Plants inhabitng in arid and semi-arid ecosystems, such as *Citrullus colocynthis* (L.) Schrad., are adapted to drought, heat, salinity, and nutrient limitation. Their associated microbial communities may further support plant persistence under these harsh conditions. Here, we characterized the bacterial communities associated with leaf endosphere, rhizosphere and roots of *C. colocynthis* growing in a semi-arid region of Moroccan using 16S rRNA gene amplicon sequencing, culture-dependent isolation, and genome-informed functional profiling of selected isolates. The results revealed a structured microbiome, with rhizosphere harboring the highest bacterial diversity, roots representing an intermediate selective habitat, and the leaf endosphere containing a more restricted assemblage. Communities were dominated by members of the phyla Pseudomonadota, Actinomycetota, Bacillota, and Bacteroidota. Several families associated to plant colonization, nutrient mobilization, and stress tolerance, including Pseudomonadaceae, Microbacteriaceae, Rhizobiaceae, Devosiaceae, and Xanthomonadaceae, showed compartment-specific enrichment. Although soil physicochemical properties influenced bacterial community structure, they explained only part of the variation observed, suggesting that bacteriome assembly is shaped by both environmental conditions and host filtering processes. Culture-bdependant analyses recovered diverse endophytic genera, mainly *Achromobacter, Pseudomonas*, and *Glutamicibacter,* most of which were also detected in the amplicon sequencing dataset. Genome-based profiling identified traits related to stress response, osmoprotection, nutrient-related metabolism, colonization, and plant–microbe interactions. Together, these findings highlight *C. colocynthis* as a reservoir for functionally relevant bacterial diversity with ecological and biotechnological potential in semi-arid environments.

**Importance:** Understanding how plants survive in arid and semi-arid ecosystems is increasingly important in the context of climate change and land degradation. This study demonstrates that *Citrullus colocynthis* hosts a structured and functionally diverse bacteriome across the leaf endosphere, rhizosphere, and root compartments. By combining amplicon sequencing, cultivation, and genome-informed functional analyses, we identified bacterial taxa and traits associated with stress tolerance, nutrient acquisition, and plant colonization. The recovery of cultivable endophytes with adaptive genomic features highlights the potential of desert plant-associated microbiota as a source of beneficial microorganisms for sustainable agriculture and biotechnological applications in water-limited environments.

---

Plants growing in arid and semi-arid environments are exposed to multiple environmental constraints, including drought, salinity, high temperatures, and nutrient limitation (1). In these ecosystems, survival depends on a tightly integrated combination of morphological, physiological, genetic, ecological, and microbial adaptations that collectively regulate water acquisition, stress tolerance, and reproductive success (2–4). Increasingly, plant performance in drylands is understood not only as a function of host traits, but as an emergent property of the plant–microbe holobiont, where root-associated microbial communities extend the adaptive capacity of plants under environmental stress (5–7).

At the structural level, hot desert plants exhibit extensive morphological and anatomical adaptations such as deep or highly branched root systems, modified xylem architecture, reduced leaf area, thick cuticles, and succulent tissues, all of which reduce water loss and improve hydraulic efficiency (2, 3, 8–10). These traits are complemented by physiological strategies including increased water-use efficiency, stomatal regulation, osmotic adjustment through compatible solutes, and enhanced antioxidant defenses that mitigate oxidative stress induced by drought and heat (11–14). At the molecular level, selection on genes involved in water transport, stress signaling, and metabolic regulation further contributes to adaptation in extreme environments (2, 9).

In parallel, root-associated microbial communities play a central role in plant adaptation to arid conditions. Plant growth-promoting rhizobacteria (PGPR) and arbuscular mycorrhizal (AM) fungi enhance nutrient acquisition, improve soil structure, modulate phytohormone signaling, and increase drought tolerance through improved water uptake and stress mitigation (7, 8, 15, 16). Microbial production of extracellular polymeric substances (EPS) contributes to soil aggregation and water retention, while microbial regulation of hormonal pathways such as abscisic acid and auxin integrates environmental signals into plant developmental and stress-response programs (17, 18).

A key concept emerging in dryland plant–microbe ecology is the rhizosheath, defined as the layer of soil tightly adhering to roots and forming a functional interface between the plant root system and the surrounding soil environment (19–22). This rhizosphere–endosphere continuum acts as a dynamic protective barrier that enhances plant resilience to abiotic stress by stabilizing the root-zone microenvironment, reducing the formation of air gaps at the root–soil interface, and improving hydraulic continuity and water transport under drought conditions. The rhizosheath also regulates nutrient fluxes, contributes to soil aggregation, and suppresses soil-borne pathogens, thereby supporting plant performance in resource-limited ecosystems. Importantly, it constitutes a favorable niche for drought-adapted and plant growth-promoting microorganisms, including rhizobacteria and AM fungi, that facilitate water and nutrient acquisition and modulate plant stress responses. These microbial communities are often enriched in taxa capable of producing EPS, osmoprotectants, phytohormones, and other stress-alleviating metabolites, collectively extending the physiological tolerance of plants under extreme environmental conditions (19, 21, 22).

*Citrullus colocynthis* (L.) Schrad., commonly known as bitter apple or desert gourd, is a wild cucurbit species widely distributed across arid and semi-arid regions of Africa, the Middle East, and South Asia. It thrives in deserts, dry riverbeds, sandy soils, and degraded lands (23, 24). Its ecological success is associated with a deep tuberous root system, efficient water acquisition and storage, reduced transpiration, osmoprotectant accumulation, and high genetic diversity (25, 26). In addition to its ecological importance, *C. colocynthis* is widely studied for its phytochemical richness, including alkaloids, flavonoids, glycosides, phenolics, and terpenoids, which contribute to antimicrobial, antioxidant, antidiabetic, anti-inflammatory, and anticancer activities (23, 24, 27). However, despite extensive phytochemical characterization, the microbial communities associated with this species remain poorly understood, particularly in relation to their ecological roles in stress adaptation.

Previous studies have reported diverse endophytic and rhizospheric bacteria associated with *C. colocynthis*, including taxa with plant growth-promoting and antimicrobial properties (28). Amplicon sequencing-based surveys further indicate that its bacterial community includes diverse soil- and plant-associated taxa linked to nutrient cycling and stress resilience (29). More broadly, plant microbiome research shows that microbial community assembly is shaped by host compartment, soil properties, environmental conditions, and host genotype, and that these communities are increasingly recognized as key contributors to plant adaptation in stressed ecosystems (30–32). However, microbiome structure and function remain highly underexplored for *C. colocynthis* across different geographic regions.

Despite growing recognition of the importance of plant–microbe interactions in dryland ecosystems, several key knowledge gaps remain. First, the microbiome of *Citrullus colocynthis* has not been comprehensively characterized across plant compartments (foliar tissues, roots, and rhizospheric soil), particularly in North African semi-arid environments. Second, the extent to which cultivable endophytes reflect the broader bacterial community remains poorly understood. Third, the functional potential of microbial communities associated with this drought-adapted species has not been investigated through an integrated community- and genome-based framework.

This study aimed to characterize the bacterial communities associated with *C. colocynthis* in a semi-arid Moroccan ecosystem and to link bacteriome composition with cultivable diversity and genome-resolved functional traits. We hypothesized that (i) bacterial communities differ among foliar, root, and rhizospheric soil compartments due to strong microenvironmental filtering; (ii) root-associated and rhizospheric bacteriomes are enriched in taxa involved in plant growth promotion and drought adaptation, consistent with the concept of a functional rhizosheath; and (iii) the cultivable fraction of the bacteriome represents dominant taxa detected by amplicon sequencing and harbors genomic traits associated with plant colonization and environmental stress tolerance.

## Materials and Methods

### Sample collection and processing

Samples were collected near the Green City of Benguerir, Morocco, from a dry wastewater drainage channel where *Citrullus colocynthis* was naturally growing (32°11′48.6″N, 7°56′30.0″W). Two sampling campaigns were performed. The first campaign was conducted in June 2022 and included 35 plants from five populations, while the second was conducted in July 2023 and included 30 plants from six populations. The sampled populations were distributed along the drainage channel, and sampling intensity varied according to population size, with 3 to 20 plants collected per population to avoid overharvesting. For each plant, root and shoot tissues were collected together with rhizospheric soil. Samples were placed in sterile plastic bags, transported to the laboratory on ice packs, and prepared upon arrival.

Plant samples were washed under running water to remove soil and debris. Root and foliar tissues were separated using sterile blades, surface-disinfected with 70% ethanol, rinsed with sterile distilled water, and dried on sterile paper towels. Plant tissues and sieved soil samples from both sampling campaigns were stored at −20°C until DNA extraction for 16S rRNA gene amplicon sequencing.

Bacterial endophytes were isolated from root and foliar tissues collected during the first sampling campaign to obtain viable isolates for culture-based characterization. Isolation, purification, and preservation followed procedures described previously (33). Briefly, surface-sterilized tissue fragments were placed on culture media, and emerging colonies with distinct morphologies were purified by repeated subculturing. Purified isolates were stored at −80°C in glycerol stocks. In parallel, genomic DNA was extracted from fresh bacterial biomass according to the protocol reported previously (34) and stored at −20°C.

### Soil physicochemical analysis

Composite soil samples were prepared for each subregion and analyzed at the AITTC laboratory, University Mohammed VI Polytechnic, Benguerir, Morocco. A total of 20 physicochemical parameters were measured, including granulometry, pH, electrical conductivity, nitrate, ammonium, total nitrogen, P₂O₅, K₂O, CaO, CaCO₃, Fe, Zn, Cu, Mn, Na₂O, MgO, organic matter, and cation exchange capacity.

### 16S rRNA gene sequencing and phylogenetic analysis of cultured isolates

Preliminary taxonomic identification of cultured bacterial isolates was performed using 16S rRNA gene Sanger sequencing, following the procedure described previously (34). Forward and reverse sequence reads were processed using Geneious Prime v2023.0.4 (Biomatters, New Zeland). Low quality terminal regions were trimmed, and forward and reverse seqeuences from each isolate were aligned, inspected, and assembled into curated consensus 16S rRNA gene sequences. Sequences marked as noise, low quality, or unsuitable for alignment were excluded from downstream analysis. The retained sequences were organized in FASTA format.

Taxonomic assignment of bacterial isolates was performed using local BLAST nucleotide searches against a 16S rRNA gene reference database (35) with BLAST+ (36). Closest reference matches were identified based on percentage identity, query coverage, E-value, and bit score. Curated 16S rRNA gene sequences were subsequently deposited in GenBank, with the final dataset comprising 112 sequences.

A complete 16S rRNA gene phylogenetic tree was first generated using all GenBank-submitted cultured isolate sequences together with their closest reference sequences. Sequences were aligned using MAFFT v7.526 (37), and phylogenetic inference was performed using FastTree v2.2.0 (38) under a nucleotide substitution model appropriate for 16S rRNA gene sequences. Branch support values were displayed at internal nodes, and tree visualization and annotation were performed using iTOL (39). Reference sequences were included to provide phylogenetic context for the taxonomic placement of the cultured isolates.

To provide a simplified overview of the cultured collection, a representative 16S rRNA gene phylogeny was additionally generated using the same alignment and tree-building workflow. For this representative tree, isolate sequences were summarized at the genus level and analyzed together with closely related reference sequences, allowing the main taxonomic groups recovered from Citrullus colocynthis to be visualized more clearly.

### DNA extraction and 16S rRNA gene amplification

Total genomic DNA was extracted from soil, root, and shoot samples from both sampling campaigns. For soil samples, 250 mg of material was processed using the DNeasy PowerSoil Pro Kit, whereas 50 mg of root or shoot tissue was extracted using the DNeasy Plant Pro Kit, following the manufacturer’s instructions (Qiagen, Global Diagnostic Distribution, Témara, Morocco). Before extraction, samples were homogenized using a TissueLyser II with 2 mm tungsten beads (Giagen) at 24 Hz for 15 min. DNA quality and quantity were assessed by agarose gel electrophoresis and spectrophotometric measurement using a BioSpectrophotometer (Eppendorff, Humborg, Germany).

The V5–V6 region of the bacterial 16S rRNA gene was amplified using HPLC purified primers containing custom sequence adapters at the 5′ end: CS1-719F 5′-ACACTGACGACATGGTTCTACA-AACMGGATTAGATACCCKG-3′ and CS2-1115R 5′-TACGGTAGCAGAGACTTGGTCT-AGGGTTGCGCTCGTTG-3′ (40). Each 25 µL PCR reaction contained 1× Platinum Direct PCR Universal Master Mix (Thermo Fisher Scientific, Rabat Morocco), 0.2 µM of each primer, and approximately 10 ng of template DNA. Amplifications were performed in duplicate using a Mastercycler X50s (Eppendorff, Hamburg, Germany) under the following conditions: initial denaturation at 94°C for 3 min; 35 cycles of 94°C for 30 s, 55°C for 30 s, and 72°C for 1 min; followed by a final extension at 72°C for 5 min. No-template controls containing sterile Milli-Q water were included in each PCR run.

### Amplicon library preparation and sequencing

Amplicon library preparation was performed following the protocols described by (21, 41). PCR products were purified using Agencourt AMPure XP beads, washed twice with ethanol, air-dried, and resuspended in 10 mM Tris buffer at pH 8.5. A second PCR was performed to attach Illumina sequencing adapters and index tags. Indexing reactions contained 5 µL of purified PCR product, 2.5 µL of Fluidigm Access Array Barcode 384, and 1× KAPA HiFi HotStart ReadyMix in a final volume of 50 µL. The indexing PCR consisted of an initial denaturation at 95°C for 3 min, followed by eight cycles of 95°C for 30 s, 55°C for 30 s, and 72°C for 30 s, with a final extension at 72°C for 5 min.

Indexed amplicons were purified using Agencourt AMPure XP beads and quantified using a Qubit fluorometer with the DNA HS kit. Library quantification, normalization, and pooling were performed according to Illumina recommendations. The final 16S rRNA gene amplicon libraries were sequenced on an Illumina MiSeq platform using a MiSeq Reagent Kit v3 with paired-end sequencing.

### 16S rRNA gene amplicon processing and ASV inference

Paired-end 16S rRNA gene amplicon reads were processed in R v4.4.2 using DADA2 v1.34.0 following the standard DADA2 workflow (42). Raw read quality profiles were inspected, and primer/adaptor sequences were removed using cutadapt v5.2 prior to denoising (43). Reads were then quality filtered, denoised, merged, and chimera-filtered using DADA2 to generate a non-chimeric amplicon sequence variant (ASV) table for downstream community analyses.

Taxonomic assignment was performed using an RDP-based taxonomy workflow (44, 45). Non-prokaryotic and off-target assignments, including plant-derived, chloroplast, mitochondrial, eukaryotic, fungal, and metazoan sequences, were removed, while archaeal ASVs were retained. The final filtered prokaryotic ASV table was combined with sample metadata in phyloseq v1.54.0 and used for downstream analyses (46).

### Amplicon sequencing community analyses

Taxonomic composition was summarized from the filtered prokaryotic ASV table at the phylum, family, and genus levels. For each taxonomic level, only ASVs assigned at that level were retained, and relative abundance was calculated among assigned taxa. Mean relative abundance was then calculated for each compartment–year group. Dominant assigned taxa were visualized as stacked bar plots, while remaining assigned taxa were grouped as “Other”. Data processing was performed in R v4.4.2 (47) using dplyr v1.2.1 (https://dplyr.tidyverse.org/), tidyr v1.3.2 (https://tidyr.tidyverse.org/), and readr v2.2.0 (https://readr.tidyverse.org/). Stacked bar plots were generated using ggplot2 v4.0.3 (https://ggplot2-book.org/), with percentage axis formatting performed using scales v1.4.0 (https://scales.r-lib.org/).

Alpha diversity was calculated from the curated non-chimeric ASV table using the Shannon and inverse Simpson indices (48). Because samples from the two sampling campaigns were stored for different durations before DNA extraction, the year grouping was not treated as a purely temporal biological factor, as storage conditions and pre-extraction handling can affect inferred microbial community profiles in amplicon sequencing studies (49, 50). Accordingly, alpha-diversity comparisons were performed among compartments within each year group. Due to non-normal distributions and unequal variances, differences in alpha diversity among compartments were assessed using Kruskal–Wallis tests followed by pairwise Wilcoxon rank-sum tests with Holm correction (51–53). Alpha-diversity distributions were visualized as boxplots.

Beta diversity was assessed using Bray–Curtis dissimilarity after transforming ASV counts to relative abundance (54). Community structure was visualized by principal coordinates analysis. Compartment effects were tested separately within each year group using PERMANOVA with 999 permutations (55). Homogeneity of multivariate dispersion was evaluated using betadisper, followed by permutation testing (53). Pairwise PERMANOVA and pairwise dispersion tests were performed within each year group, with P-values adjusted using Holm correction (53) Differential abundance analysis was performed using ANCOM-BC2 implemented in the ANCOMBC v2.12.0 package (56). The filtered ASV table, taxonomy table, and metadata were combined into a phyloseq object using phyloseq v1.54.0 (46). ANCOM-BC2 was run separately for each year group, with sample compartment as the fixed effect. Analyses were performed independently at the phylum, family, and genus levels. Pairwise comparisons were conducted for roots vs foliar, soil vs foliar, and soil vs roots. Models were run with structural-zero detection, negative lower-bound correction, and pseudo-count sensitivity analysis enabled, using Holm correction and a significance threshold of α = 0.05. Family-level results were used for the main visualization because they provided the best balance between taxonomic resolution and interpretability.

### Association between soil physicochemical properties and community structure

Associations between regional physicochemical properties and prokaryotic community structure were assessed using constrained analysis of principal coordinates/distance-based redundancy analysis based on Bray–Curtis dissimilarities (57). Analyses were conducted in R using phyloseq and vegan. ASV counts were transformed to relative abundance before Bray–Curtis dissimilarities were calculated.

Physicochemical variables were available at the year–region level and were assigned to sequencing samples according to their corresponding year and region. Models were fitted separately for each year group and included sample compartment as a conditioning factor using Condition(Origin). Reduced environmental models were defined based on ecological relevance and correlation screening to avoid overparameterization. Overall model significance and individual environmental terms were assessed using permutation tests with 999 permutations, and model explanatory power was summarized using R² and adjusted R².

### Comparison between cultured isolates and amplicon-sequencing detected taxa

To evaluate the overlap between the cultured isolate collection and the amplicon-detected community, Sanger-based genus assignments of cultured isolates were compared with genus-level assignments in the filtered amplicon taxonomy table. For each cultured genus, detection in the amplicon dataset was recorded based on the presence of the corresponding genus in the filtered ASV table. Detection was additionally summarized by compartment, including foliar, root, and soil samples. This comparison was interpreted as genus-level taxonomic correspondence only and does not imply detection of the same cultured isolate or strain in the amplicon dataset.

### Genome-level functional characterization of selected cultured isolates

Whole-genome sequence data were used to provide functional context for selected cultured isolates. Sample preparation, sequencing, read preprocessing, assembly, and initial taxonomic analyses of the genome-sequenced isolates were performed as described previously (33). Briefly, raw reads were quality filtered using a Phred quality threshold of Q20 and a minimum read length of 100 bp, assembled de novo, and evaluated using standard genome-quality metrics including completeness, contamination, N50, contig number, genome size, and GC content.

For the present study, redundant genome assemblies corresponding to the same species-level clusters were removed before functional summarization, retaining one genome per redundant cluster based on taxonomic relatedness and assembly quality. The final non-redundant genome set included 17 selected cultured isolates. Genome annotation was performed using Prokka v1.14.6 (58) with the bacterial kingdom setting. Prokka annotation tables were screened using custom Python 3 scripts to identify curated functional marker sets associated with plant–microbe interactions and environmental adaptation, including ACC deaminase, siderophore/iron acquisition, nitrogen-related metabolism, osmoprotection, oxidative stress response, general stress response, motility/chemotaxis, biofilm/adhesion, secretion systems, and antagonism-associated markers.

For each functional system, predefined marker groups were searched in Prokka gene names and product annotations using regular-expression matching. Marker-set completeness was calculated as the percentage of predefined marker groups detected in each genome. Data processing and reshaping were performed in R using dplyr, tidyr, and readr, and the resulting matrix was visualized as a heatmap using ggplot2.

## Results

### 16S rRNA gene sequencing output and ASV dataset quality

The curated DADA2 tracking table included 174 samples. Across all samples, 19,358,967 input reads were processed, of which 18,502,284 reads were retained after quality filtering, corresponding to a quality-filtering retention of 95.57%. After denoising, paired-end read merging, and chimera removal, the final non-chimeric ASV table contained 14,528,658 reads, representing 75.05% of the input reads (Table S1). The average number of reads per sample was approximately 83,500.

### Taxonomic composition of the bacterial community

Community composition was summarized at the phylum, family, and genus levels across compartments and year groups. At the phylum level, Pseudomonadota represented a major component of the assigned community in plant-associated compartments, particularly in leaf endosphere and root samples. Rhizosphere samples showed a distinct broader taxonomic profile, with greater representation of Chloroflexota and other phyla. Differences in the relative contribution of dominant phyla were observed across compartment and year groups (Figure S1). At the family level, dominant assigned families differed among compartments. Pseudomonadaceae predominated in plant-associated samples, whereas Tepidiformaceae was more abundant in rhizosphere-associated communities. Several other families, including Bacillaceae, Erwiniaceae, Nocardioidaceae, Trueperaceae, and Micrococcaceae, exhibited variable relative abundances across compartments and sampling years. A substantial proportion of sequences was assigned to the “Other” category, reflecting the contribution of numerous low-abundance families to the overall community structure (Figure 1).

**Figure 1.**
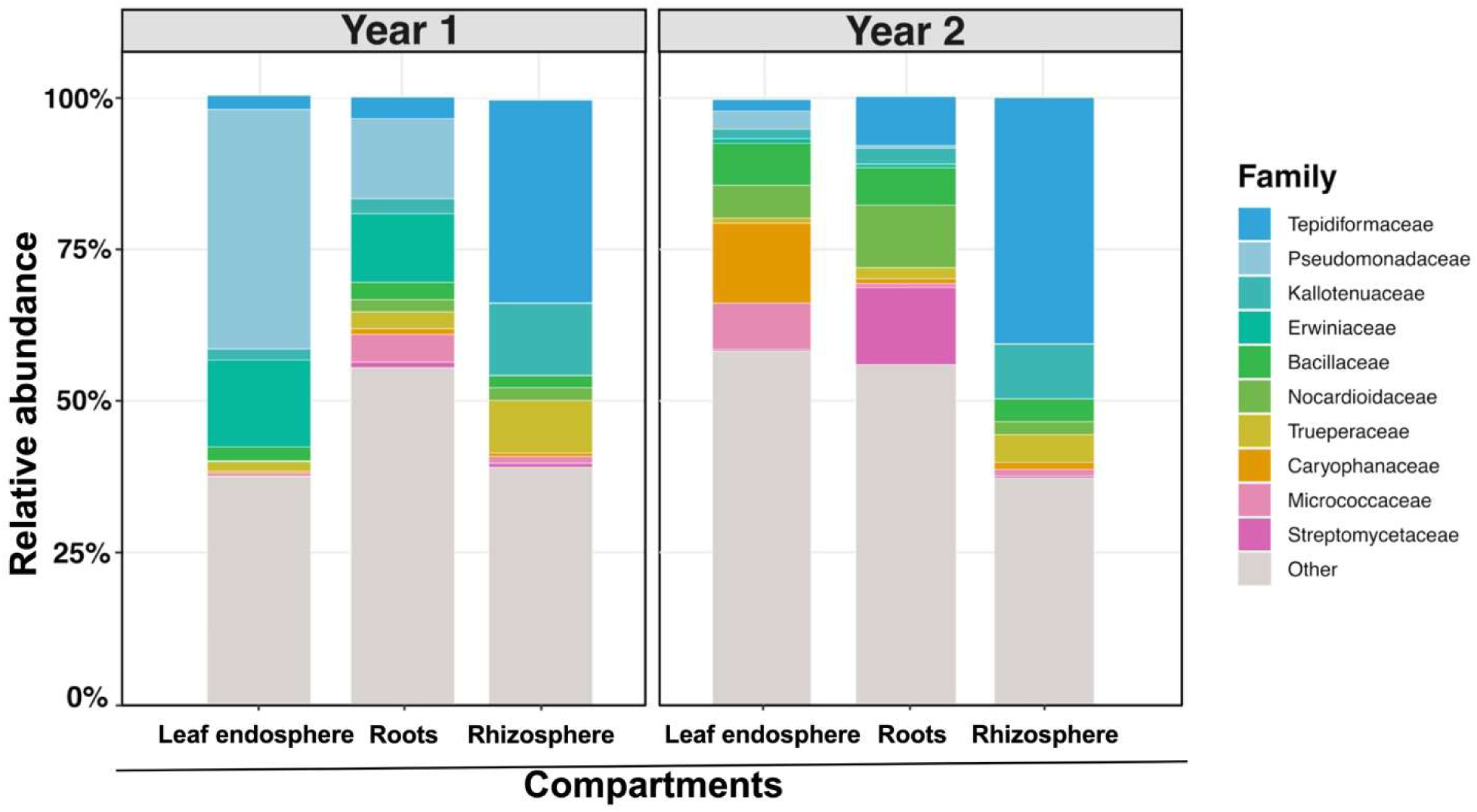
Family-level composition of bacterial communities across compartments and years. Stacked bar plots show the mean relative abundance of the dominant bacterial families in leaf endosphere, root, and rhizosphere samples collected during Year 1 and Year 2. Families not included among the dominant taxa were grouped as “Other.” Relative abundances were calculated from ASVs assigned at the family level. Colors indicate bacterial families.

At the genus level, community composition was more heterogeneous, with the “Other” category accounting for a substantial proportion of assigned genera across several compartment–year combinations. *Pseudomonas* was among the most abundant genera in plant-associated compartments, whereas genera such as *Nocardioides*, *Truepera*, *Kaistella*, *Brevundimonas*, and *Tepidiforma* showed variable distributions among compartments and sampling years. Overall, the taxonomic profiles revealed clear compartment-specific patterns in bacterial community composition, with temporal variation becoming more pronounced at finer taxonomic resolution (Figure S2).

### Alpha diversity differs among compartments

Alpha diversity varied significantly among compartments within both year groups. Shannon diversity differed among leaf endosphere, root, and rhizosphere samples in Year 1 (Kruskal–Wallis χ² = 59.70, df = 2, p = 1.09 × 10⁻¹³) and Year 2 (χ² = 55.07, df = 2, p = 1.10 × 10⁻¹²) (Figure 2, Table S2). The same pattern was observed for inverse Simpson diversity in Year 1 (χ² = 60.02, df = 2, p = 9.28 × 10⁻¹⁴) and Year 2 (χ² = 43.13, df = 2, p = 4.30 × 10⁻¹⁰) (Figure 3, Table S2). airwise Wilcoxon rank-sum tests with Holm correction showed significant differences among all compartment pairs for both alpha-diversity indices in both year groups (Table S3). Soil samples were consistently the most diverse compartment. In Year 1, foliar samples showed the lowest diversity, while root samples were intermediate. In Year 2, foliar diversity was higher than in Year 1, although this difference should be interpreted cautiously because the year groups differed in storage duration. Supporting aligned rank transform ANOVA confirmed a strong compartment effect for both Shannon and inverse Simpson diversity. For Shannon diversity, compartment, year, and their interaction were significant, whereas for inverse Simpson diversity, compartment and the compartment–year interaction were significant, while the overall year effect was not (Table S4).

**Figure 2.**
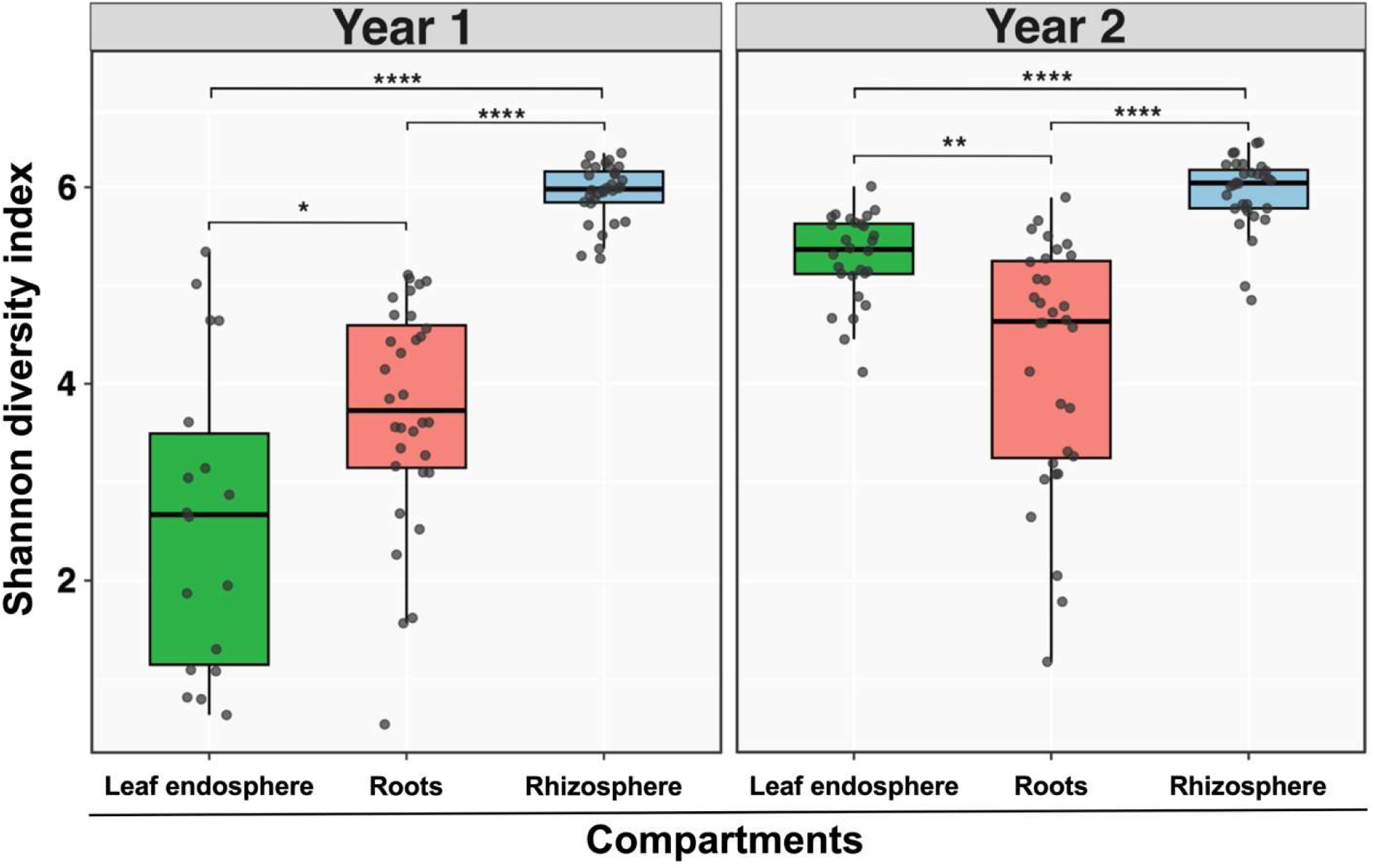
Shannon diversity across leaf endosphere, root, and rhizosphere compartments by year. Boxplots show Shannon alpha diversity of bacterial communities in leaf endosphere, root, and rhizosphere compartments, displayed separately for Year 1 and Year 2. Points represent individual samples, and brackets indicate significant pairwise differences among compartments within each year.

**Figure 3.**
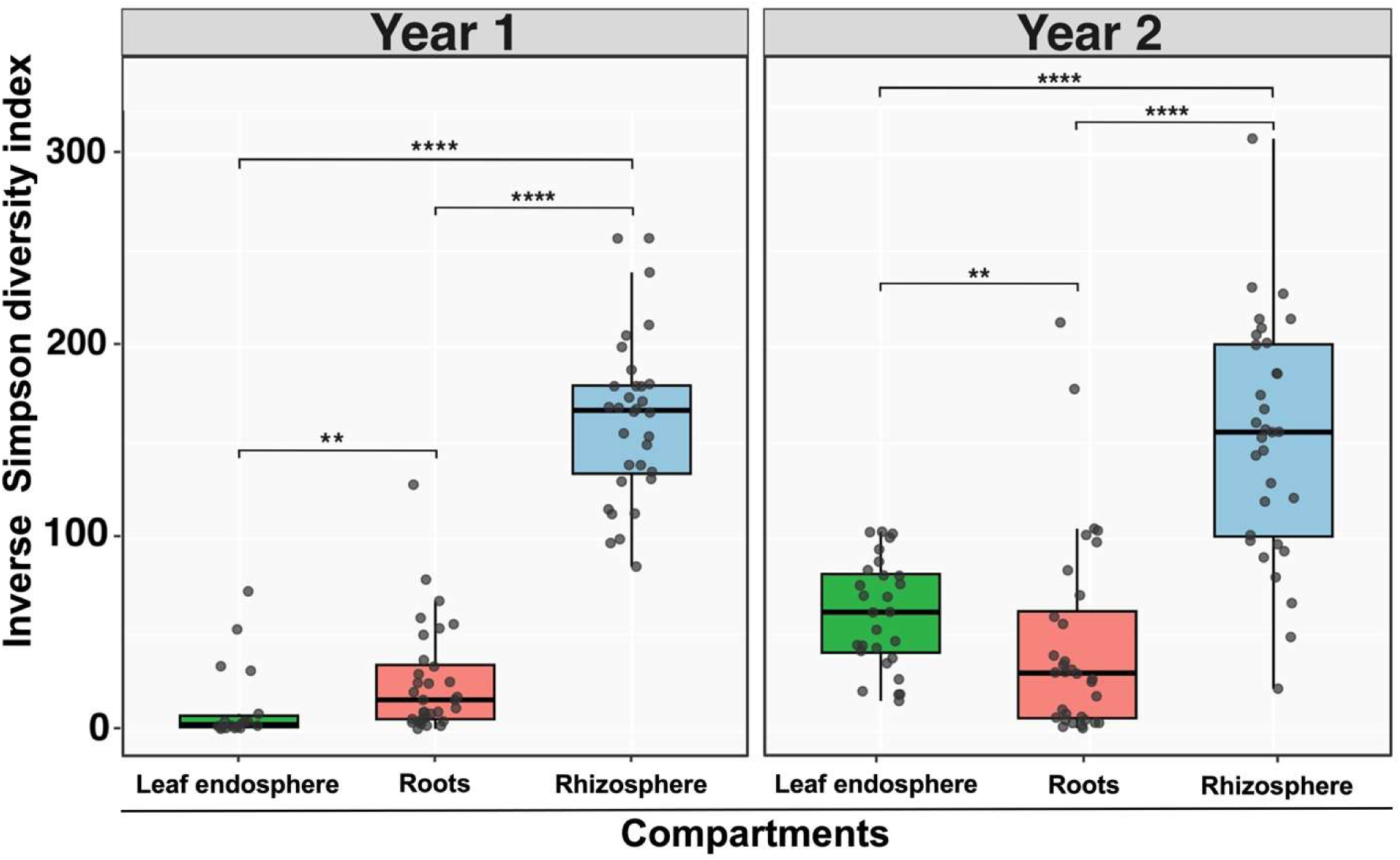
Inverse Simpson diversity across leaf endosphere, root, and rhizosphere compartments by year. Boxplots show Inverse Simpson alpha diversity of bacterial communities in leaf endosphere, root, and rhizosphere compartments, displayed separately for Year 1 and Year 2. Points represent individual samples, and brackets indicate significant pairwise differences among compartments within each year.

### Beta-diversity structure across compartments

Principal coordinates analysis based on Bray–Curtis dissimilarities showed clear compartment related structuring of bactrial community composition within both year groups (Figure 4). PERMANOVA confirmed significant differences among compartments in Year 1 (R² = 0.2803, F = 15.3860, p = 0.001) and Year 2 (R² = 0.2920, F = 18.3540, p = 0.001), indicating that compartment explained approximately 28–29% of the variation in Bray–Curtis community structure within each year group (Table S5).

**Figure 4.**
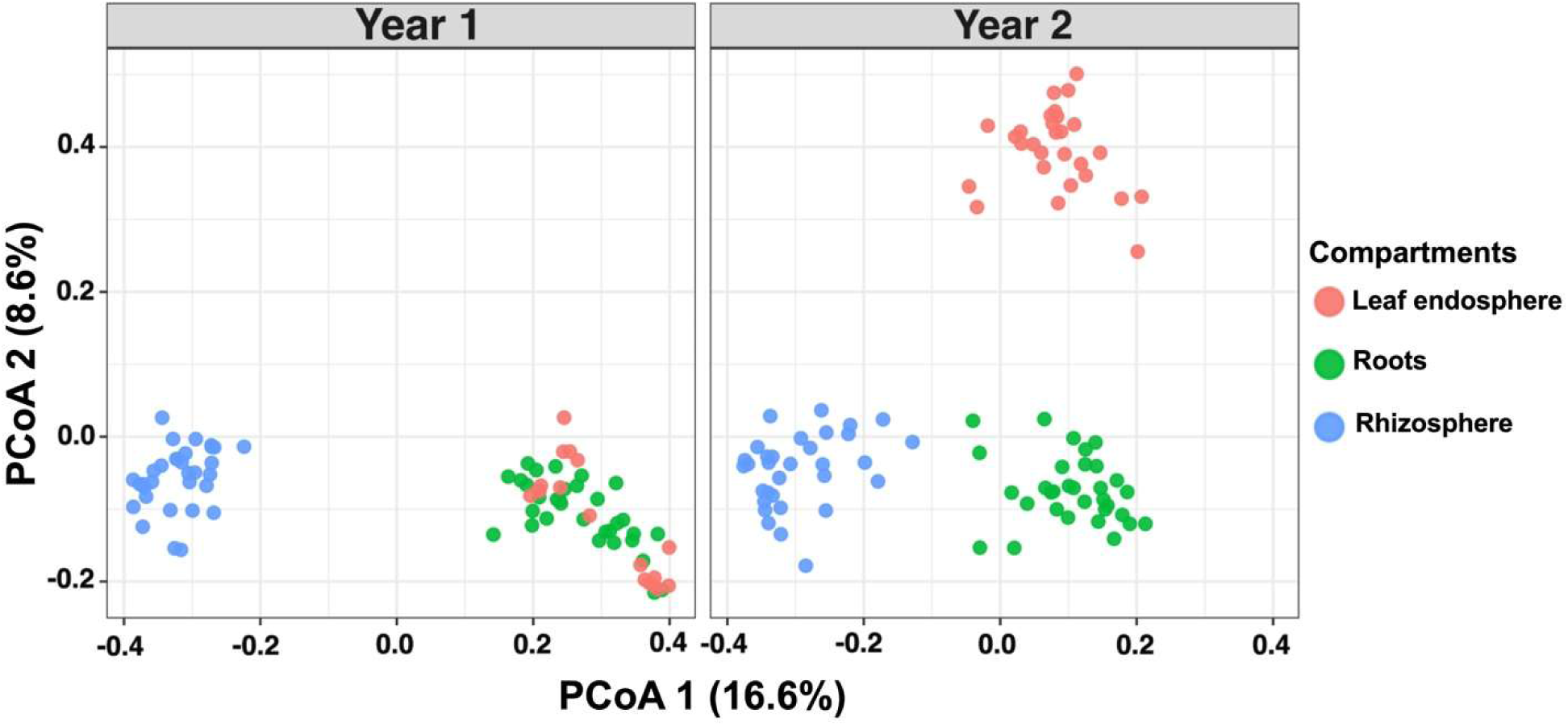
Principal coordinates analysis of Bray–Curtis dissimilarities by compartment and year. Principal coordinates analysis (PCoA) based on Bray–Curtis dissimilarities showing variation in bacterial community composition among leaf endosphere, root, and rhizosphere compartments across Year 1 and Year 2. Each point represents an individual sample and is colored according to compartment. The percentage of variance explained by each axis is indicated on the corresponding axis label.

However, homogeneity of dispersion tests were also significant in both Year 1 (F = 62.3777, p = 0.001) and Year 2 (F = 111.4881, p = 0.001), indicating that within-compartment dispersion differed among groups. Therefore, the observed beta-diversity patterns reflect both compartment-associated differences in community composition and differences in within-group heterogeneity. All pairwise compartment contrasts were significant by PERMANOVA after Holm correction in both year groups (adjusted p = 0.003 for all comparisons). Pairwise dispersion tests were also significant for all comparisons except foliar versus roots in Year 1, suggesting that several pairwise beta-diversity differences were accompanied by unequal within-group dispersion (Table S6).

### Differentially abundant families across compartments

ANCOM-BC2 was used to identify robust differentially abundant families among leaf endosphere, rhizosphere, and root compartments, with analyses performed separately for each year group. Family-level results were used for visualization because they provided a balance between taxonomic resolution and interpretability. Across pairwise compartment comparisons, the number of robust differentially abundant families ranged from 18 to 56 per contrast. For each comparison, the top 15 families ranked by absolute log-fold change were visualized (Figure 5; Table 2).

**Figure 5.**
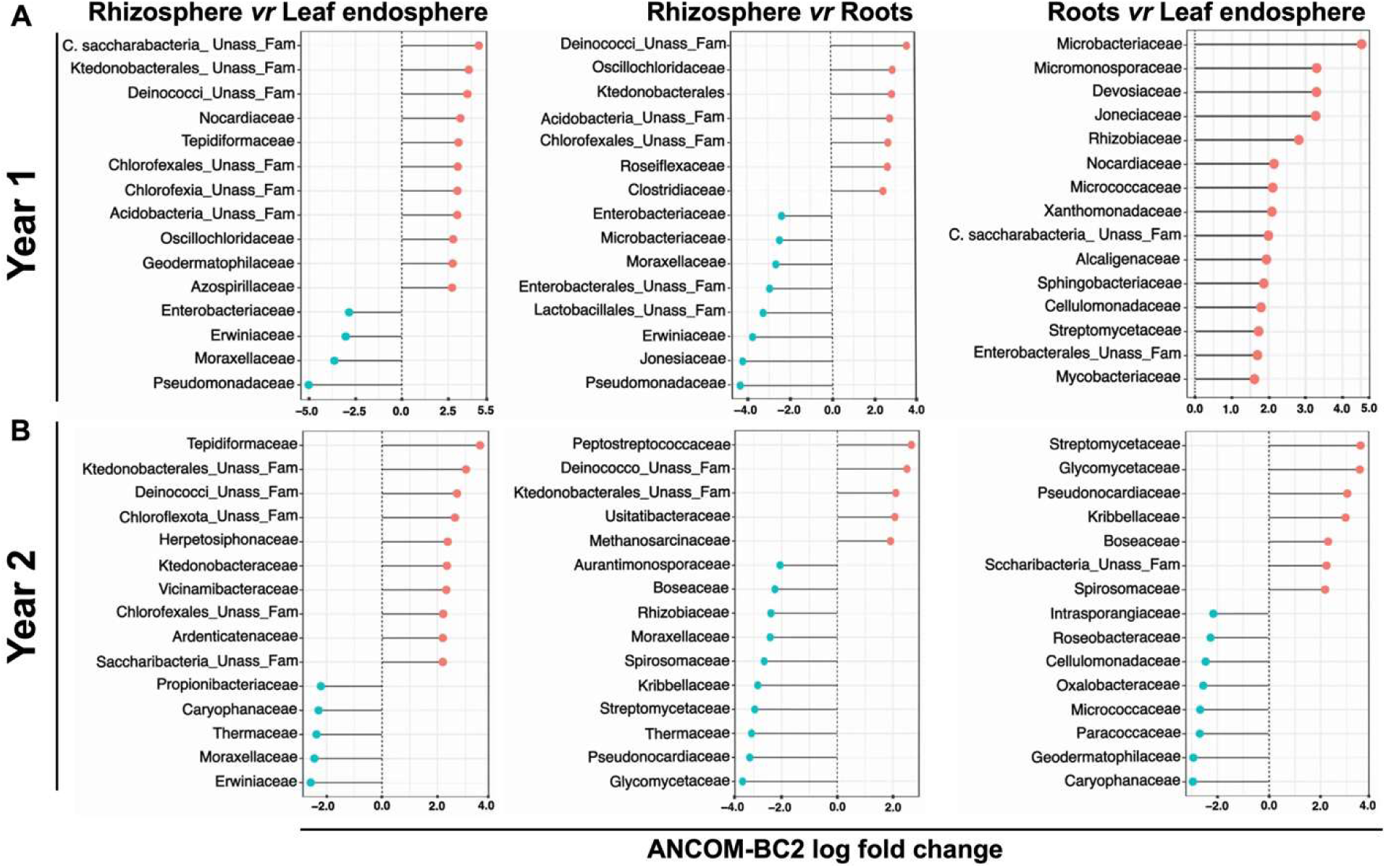
Family-level differential abundance across compartments. Top 15 robust differentially abundant bacterial families identified by ANCOM-BC2 in pairwise comparisons among leaf endosphere, root, and rhizosphere compartments in Year 1 (A) and Year 2 (B). Each panel represents a single pairwise comparison. The x-axis shows the ANCOM-BC2 log fold change, and the dashed vertical line indicates no difference between compartments. Positive values indicate enrichment in the first compartment named in the panel title, whereas negative values indicate enrichment in the second compartment. Taxa are shown at the family level; “Unass_Fam” denotes an unassigned family within the corresponding higher-level taxonomic lineage.

**Table 1.** Pairwise Wilcoxon comparisons of alpha diversity among compartments within each year group.

| Comparison | Shannon,<br>Year 1 | Shannon,<br>Year 2 | Inverse Simpson,<br>Year 1 | Inverse Simpson,<br>Year 2 |
| --- | --- | --- | --- | --- |
| Leaf endosphere vs Roots | 0.0147 | $6.90 \times 10^{-5}$ | 0.00437 | 0.00902 |
| Leaf endosphere vs Rhizosphere | $8.86 \times 10^{-13}$ | $7.08 \times 10^{-9}$ | $2.22 \times 10^{-13}$ | $6.23 \times 10^{-9}$ |
| Roots vs Rhizosphere | $3.27 \times 10^{-18}$ | $3.95 \times 10^{-13}$ | $9.82 \times 10^{-17}$ | $5.70 \times 10^{-9}$ |

**Table 2.**
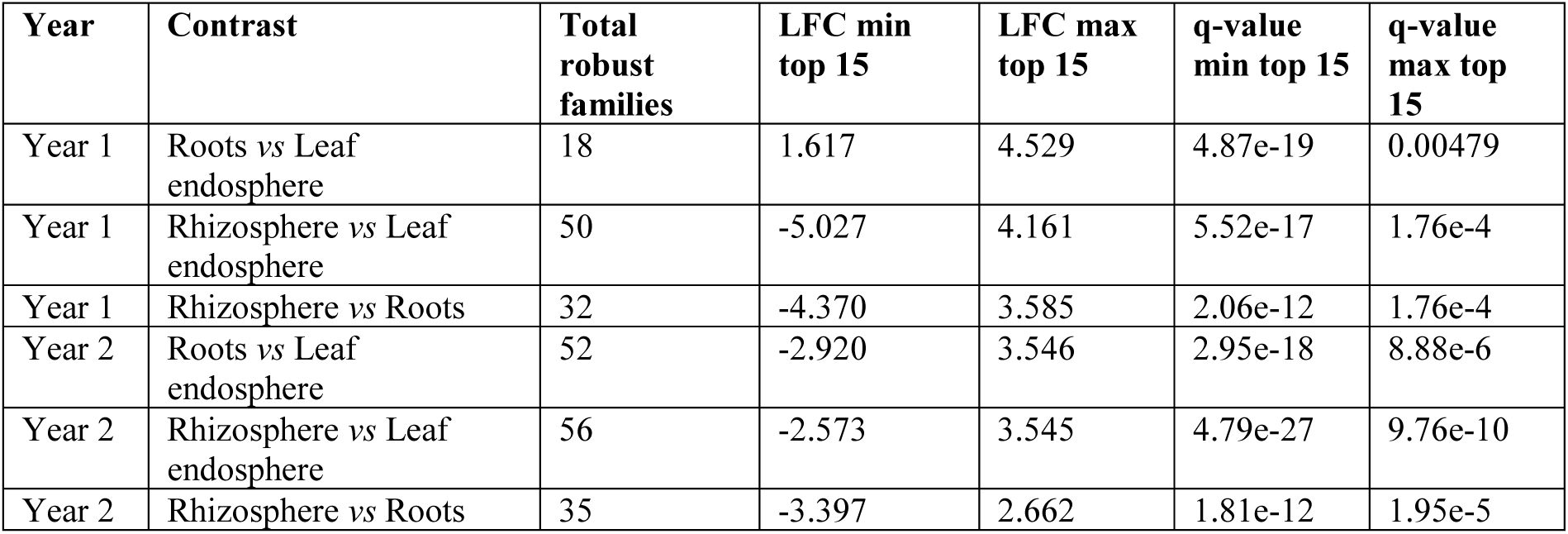
Summary statistics of family-level pairwise ANCOM-BC2 analyses. The table presents the ranges of log fold-change (LFC) values and adjusted q-values for the 15 most robust differentially abundant families identified in each pairwise comparison. Minimum and maximum LFC values correspond to the observed range among these families, while minimum and maximum adjusted q-values indicate the corresponding significance range. Results are reported separately for each year and compartment contrast.

In Year 1, the roots *versus* leaf endosphere comparison showed a root-enrichment pattern among the displayed families. Root-enriched families included several Actinomycetota and plant-associated lineages, such as Microbacteriaceae, Micromonosporaceae, Nocardioidaceae, Micrococcaceae, and Jonesiaceae, together with Rhizobiaceae, Devosiaceae, and Xanthomonadaceae. In the rhizosphere *versus* leaf endosphere comparison, rhizosphere-enriched families included environmental and soil-associated lineages such as Nocardioidaceae, Geodermatophilaceae, Myxococcaceae, Oscillochloridaceae, and unassigned families within higher soil-associated lineages. Leaf endosphere-enriched families included Pseudomonadaceae, Moraxellaceae, Enterobacteriaceae, Erwiniaceae, Staphylococcaceae, Streptococcaceae, and Enterococcaceae. The rhizosphere versus roots comparison showed enrichment in both compartments, with rhizosphere-associated families enriched on one side and root-associated families such as Microbacteriaceae, Rhizobiaceae, Devosiaceae, Pseudomonadaceae, and Xanthomonadaceae enriched on the other.

In Year 2, the rhizosphere versus leaf endosphere comparison again yielded the highest number of robust differentially abundant families, with 56 families detected. Among the top 15 displayed families, 10 were enriched in the rhizosphere and five were enriched in the leaf endosphere, with log-fold change values ranging from −2.573 to 3.545 (Table 2). The roots versus leaf endosphere comparison identified 52 robust differentially abundant families, with the top 15 families showing enrichment in both compartments: seven in roots and eight in the leaf endosphere. The rhizosphere versus roots comparison identified 35 robust differentially abundant families, with five of the top 15 families enriched in the rhizosphere and 10 enriched in roots.

Overall, the ANCOM-BC2 results at the family level showed clear compartment-associated differences within each year group. Rhizosphere versus leaf endosphere comparisons contained the highest number of robust differentially abundant families in both independently analyzed year groups, while root-associated contrasts showed distinct family-level enrichment patterns relative to the leaf endosphere and rhizosphere compartments.

### Associations between soil physicochemical properties and community structure

CAP/db-RDA was used to assess whether regional physicochemical profiles were associated with bacterial community composition after accounting for sample origin. Models were fitted separately for each year group, and origin was included as a conditioning factor to account for compartment-level effects (Table 3).

**Table 3.** Summary of CAP/db-RDA models testing associations between regional physicochemical properties and bacterial community composition.

| Model | Variables included | Year | Df | F value | p-value | R<sup>2</sup> | Adjusted R<sup>2</sup> | Significant / strongest terms |
| --- | --- | --- | --- | --- | --- | --- | --- | --- |
| Broad soil condition | pH, clay, organic matter, Olsen phosphorus, electrical conductivity | Year 1 | 5 | 1.1496 | 0.060 | 0.0523 | 0.0070 | pH |
| Broad soil condition | pH, clay, organic matter, Olsen phosphorus, electrical conductivity | Year 2 | 5 | 2.3763 | 0.001 | 0.0877 | 0.0520 | pH, clay, organic matter, Olsen phosphorus, electrical conductivity |
| Nutrient/fertility | Organic matter, Olsen phosphorus, potassium, total nitrogen | Year 1 | 4 | 1.2648 | 0.015 | 0.0459 | 0.0099 | Potassium |
| Nutrient/fertility | Organic matter, Olsen phosphorus, potassium, total nitrogen | Year 2 | 4 | 2.4428 | 0.001 | 0.0730 | 0.0441 | Organic matter, Olsen phosphorus, potassium, total nitrogen |
| Texture/mineral | Clay, sand, pH, total limestone, cation exchange capacity | Year 1 | 5 | 1.1799 | 0.040 | 0.0536 | 0.0084 | Clay |
| Texture/mineral | Clay, sand, pH, total limestone, cation exchange capacity | Year 2 | 5 | 2.3763 | 0.001 | 0.0877 | 0.0520 | Clay, sand, pH, total limestone, cation exchange capacity |

In Year 1, the broad soil condition model showed a marginal association with community structure but did not reach statistical significance under permutation testing (p = 0.060, R² = 0.0523, adjusted R² = 0.0070) (Table S7), indicating a trend toward significance. The nutrient/fertility model was significant (p = 0.015, R² = 0.0459, adjusted R² = 0.0099), with potassium showing the strongest individual marginal association among the tested terms (Table S8). The texture/mineral model was also significant overall (p = 0.040, R² = 0.0536, adjusted R² = 0.0084), with clay showing the strongest individual association (Table S9). Overall, these results indicate weak but detectable associations between regional physicochemical profiles and bacterial community structure in Year 1, with low adjusted explanatory power.

In Year 2, all three reduced environmental models were significant. The broad soil condition model was associated with community structure (p = 0.001, R² = 0.0877, adjusted R² = 0.0520), with pH, clay, organic matter, Olsen phosphorus, and electrical conductivity showing significant marginal associations (Table S7). The nutrient/fertility model was also significant (p = 0.001, R² = 0.0730, adjusted R² = 0.0441), and all included variables showed significant marginal associations (Table S8). Similarly, the texture/mineral model was significant (p = 0.001, R² = 0.0877, adjusted R² = 0.0520), with clay, sand, pH, total limestone, and cation exchange capacity all significant in marginal tests (Table S9).

Together, the CAP/db-RDA results suggest that regional physicochemical profiles were associated with bacterial community composition after accounting for compartment identity. However, adjusted R² values remained low to modest across models, indicating that these environmental variables explained only a limited fraction of total community variation.

### 16S rRNA gene-based identification of cultured bacterial isolates

A total of 112 curated 16S rRNA gene sequences from cultured bacterial isolates were retained in the final GenBank-submitted dataset. Local BLASTn comparison against the 16S rRNA reference database assigned the isolates to multiple genus-level groups. The most represented genera were *Achromobacter* (n = 26), *Pseudomonas* (n = 24), and *Glutamicibacter* (n = 15). Other recovered genera included *Erwinia* (n = 10), *Enterococcus* (n = 6), *Advenella* (n = 6), *Paenibacillus* (n = 5), *Microbacterium* (n = 3), *Brucella* (n = 3), *Desemzia* (n = 2), and *Pantoea* (n = 2). Several genera were represented by a single isolate, including *Bacillus*, *Robertmurraya*, *Carnobacterium*, *Psychrobacillus*, *Sanguibacter*, *Brevibacterium*, *Arthrobacter*, *Stenotrophomonas*, *Alcaligenes*, and *Pollutimonas*.

All sequences were deposited in GenBank, and their accession numbers are provided in Table S10. Because the sequences represented partial 16S rRNA gene fragments, taxonomic assignments were interpreted conservatively at the genus level, while species-level BLAST matches were treated as closest reference matches rather than definitive species identifications.

The representative 16S rRNA gene phylogeny summarized the taxonomic structure of the cultured collection and showed clear genus-level clustering of isolates with their closest reference sequences (Figure 6). Major clusters corresponded to the dominant cultured genera, including *Achromobacter*, *Pseudomonas*, *Glutamicibacter*, *Erwinia*, *Enterococcus*, *Paenibacillus*, and *Microbacterium*. The complete isolate-level tree, including all GenBank-submitted sequences and closest reference sequences, is provided as Supplementary Figure S3.

**Figure 6.**
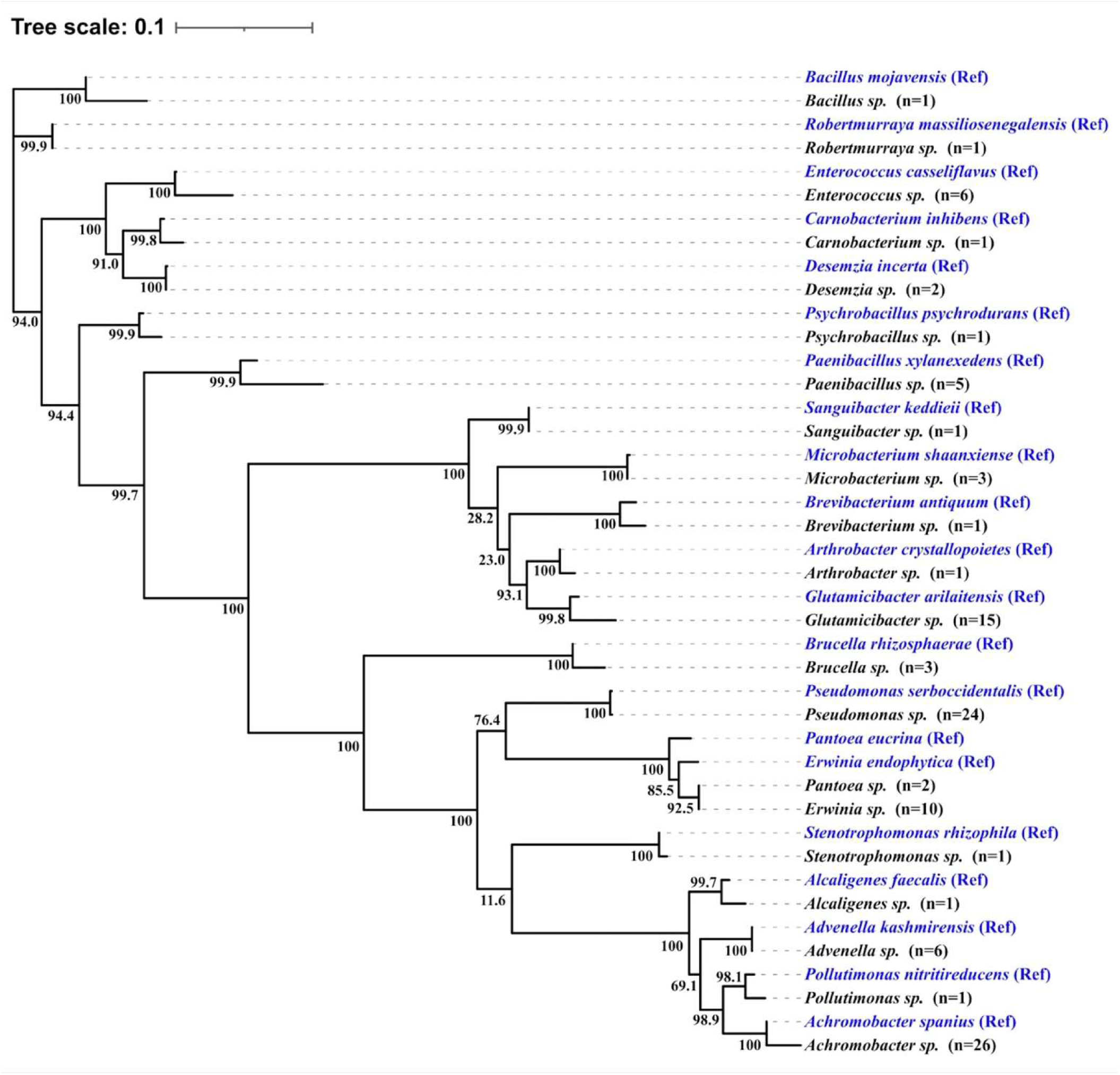
Representative 16S rRNA gene phylogeny of bacterial isolates associated with *Citrullus colocynthis*. The phylogenetic tree illustrates the taxonomic diversity of the cultured bacterial collection recovered in this study. Reference sequences are indicated by “Ref” and shown in blue, whereas isolates from this study are shown in black and grouped at the genus level, with the number of isolates represented in parentheses. Bootstrap support values (%) are displayed at internal nodes. The scale bar corresponds to 0.1 nucleotide substitutions per site, indicating the estimated evolutionary distance (sequence divergence) among taxa..

### Overlap between cultured isolates and amplicon-detected taxa

The cultured isolate collection was compared with the bacterial amplicon dataset to evaluate whether genera recovered by cultivation were also detected by amplicon sequencing. Overlap was assessed at the genus level by comparing Sanger-based genus assignments of cultured isolates with genus-level assignments in the filtered amplicon ASV table.

Most cultured genera were also detected in the amplicon dataset. Of the 21 cultured genera, 20 were detected by amplicon sequencing. Several cultured genera, including *Achromobacter*, *Advenella*, *Pollutimonas*, *Pseudomonas*, *Arthrobacter*, *Glutamicibacter*, *Erwinia*, *Pantoea*, *Enterococcus*, *Paenibacillus*, *Bacillus*, *Psychrobacillus*, *Robertmurraya*, *Brucella*, *Carnobacterium*, *Microbacterium*, *Brevibacterium*, and *Sanguibacter*, were detected across foliar, root, and soil amplicon profiles. *Alcaligenes* was detected in foliar and root samples, while *Desemzia* was detected only in root samples. *Stenotrophomonas* was the only cultured genus not detected in the filtered amplicon dataset (Table 4).

**Table 4.** Genus-level overlap between cultured isolates and the 16S rRNA gene amplicon sequening dataset.

| Cultured genera | Isolation origin | Detected in amplicon dataset | Amplicon compartments detected |
| --- | --- | --- | --- |
| *Achromobacter* | Leaf endosphere; Roots | Yes | Leaf endosphere; Roots; Rhizosphere |
| *Advenella* | Leaf endosphere; Roots | Yes | Leaf endosphere; Roots; Rhizosphere |
| *Alcaligenes* | Leaf endosphere | Yes | Leaf endosphere; Roots |
| *Pollutimonas* | Roots | Yes | Leaf endosphere; Roots; Rhizosphere |
| *Pseudomonas* | Leaf endosphere; Roots | Yes | Leaf endosphere; Roots; Rhizosphere |
| *Arthrobacter*, | Roots | Yes | Leaf endosphere; Roots; Rhizosphere |
| *Glutamicibacter* | Leaf endosphere; Roots | Yes | Leaf endosphere; Roots; Rhizosphere |
| *Erwinia*, | Leaf endosphere; Roots | Yes | Leaf endosphere; Roots; Rhizosphere |
| *Pantoea* | Leaf endosphere | Yes | Leaf endosphere; Roots; Rhizosphere |
| *Enterococcus* | Roots | Yes | Leaf endosphere; Roots; Rhizosphere |
| *Paenibacillus* | Leaf endosphere; Roots | Yes | Leaf endosphere; Roots; Rhizosphere |
| *Bacillus* | Leaf endosphere | Yes | Leaf endosphere; Roots; Rhizosphere |
| *Psychrobacillus* | Roots | Yes | Leaf endosphere; Roots; Rhizosphere |
| *Robertmurraya* | Leaf endosphere | Yes | Leaf endosphere; Roots; Rhizosphere |
| *Brucella* | Roots | Yes | Leaf endosphere; Roots; Rhizosphere |
| *Carnobacterium* | Leaf endosphere | Yes | Leaf endosphere; Roots; Rhizosphere |
| *Desemzia* | Roots | Yes | Roots |
| *Microbacterium* | Leaf endosphere; Roots | Yes | Leaf endosphere; Roots; Rhizosphere |
| *Brevibacterium* | Roots | Yes | Leaf endosphere; Roots; Rhizosphere |
| *Sanguibacter* | Leaf endosphere | Yes | Leaf endosphere; Roots; Rhizosphere |
| *Stenotrophomonas* | Roots | No | Not detected |

These results indicate broad genus level overlap. However, it only indicates taxonomic correspondence and not evidence that the same cultured isolates or strains were detected by amplicon sequencing.

### Genome-inferred functional traits associated with plant colonization, stress tolerance, and growth promotion

Functional marker-set profiling was performed on the selected genome-sequenced cultured isolates. The heatmap showed that all genomes contained marker groups associated with oxidative stress response, while general stress-response markers were detected at high completeness across nearly all genomes. Osmoprotection related marker sets were also broadly represented, with completeness values ranging from moderate to high across the genome set (Figure 7).

**Figure 7.**
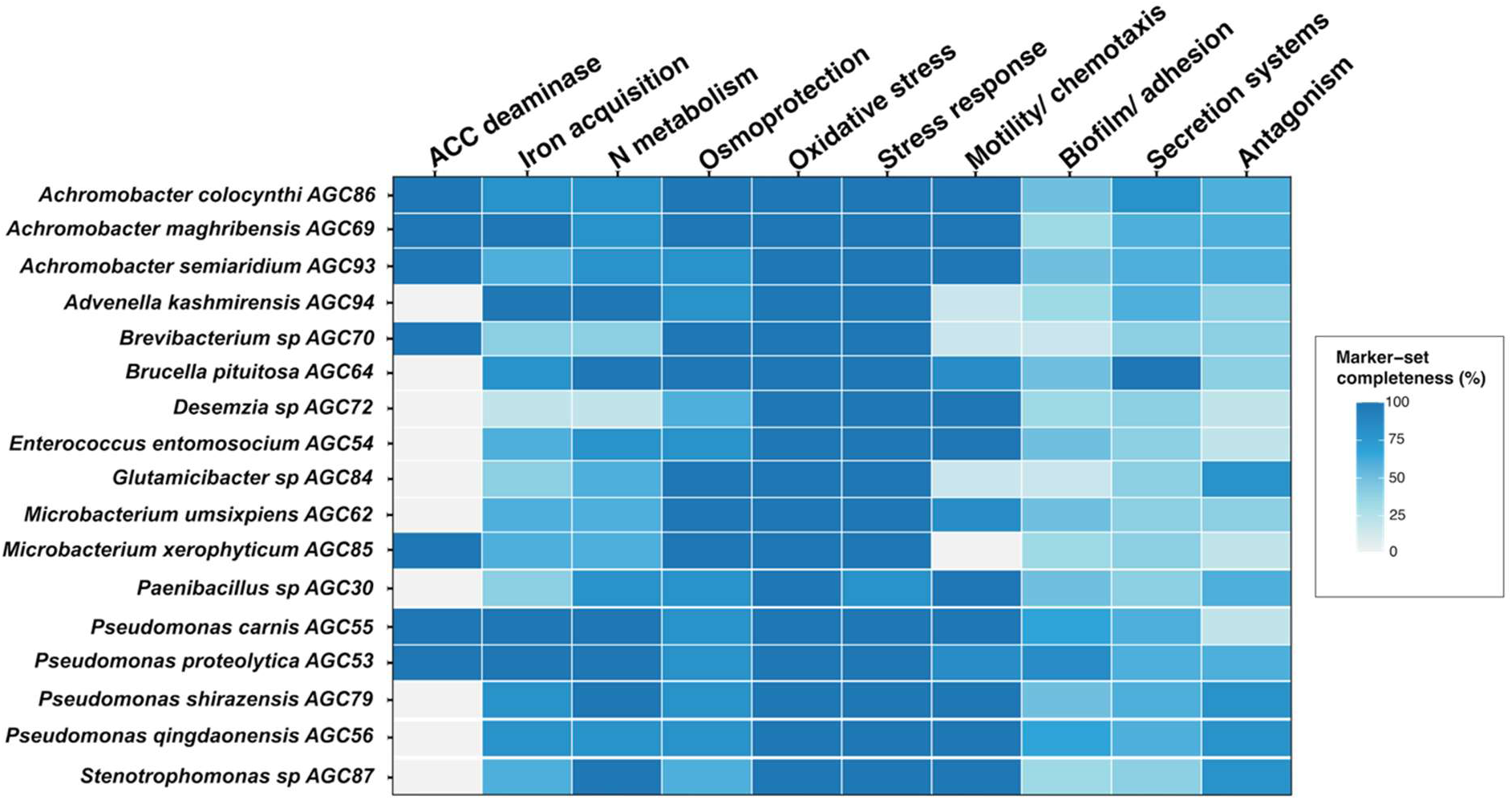
Functional marker-set completeness in genomes of selected cultured isolates. Heatmap showing the percentage completeness of curated functional marker sets across selected bacterial genomes. Functional categories include ACC deaminase, siderophore and iron acquisition, nitrogen metabolism, osmoprotection, oxidative stress response, general stress response, motility and chemotaxis, biofilm formation and adhesion, secretion systems, and antagonism-associated traits. Values represent the percentage of predefined marker groups detected in Prokka-annotated genomes, providing an overview of the functional potential of each isolate.

Siderophore/iron acquisition and nitrogen related metabolism showed more variable completeness among isolates. Siderophore/iron acquisition marker-set completeness ranged from low to complete detection, with several genomes showing high completeness, including AGC53, AGC55, AGC69, and AGC94. Nitrogen related metabolism also varied among genomes, with several isolates showing complete or near complete marker-set representation. Motility/chemotaxis showed strong variation, with some genomes showing complete marker-set detection and others showing low or no detected completeness. ACC deaminase markers were detected only in a subset of genomes, including AGC53, AGC55, AGC69, AGC70, AGC85, AGC86, and AGC93. Biofilm/adhesion, secretion systems, and antagonism-associated markers were also detected across the genome set with variable completeness.

Overall, the selected genome-sequenced cultured isolates encoded multiple functional marker sets associated with plant-associated lifestyles, with conserved detection of stress-related markers and more variable profiles for ACC deaminase, siderophore/iron acquisition, motility/chemotaxis, secretion systems, and antagonism associated markers.

## Discussion

This study provides an integrated characterization of the bacterial communities associated with *Citrullus colocynthis* in a semi-arid Moroccan environment using amplicon sequencing, culture-dependent isolation, Sanger-based identification, culture–amplicon comparison, and genome-informed functional profiling. The results suggest a structured community across the plant–soil system, in which the rhizosphere, roots, and the leaf endosphere represent distinct but interconnected microbial habitats.

Multiple complementary analyses support compartment-associated differences in community structure. Soil samples exhibited higher alpha diversity compared to plant-associated compartments. Beta-diversity analyses based on Bray–Curtis distances indicated clear separation among soil, root, and leaf endosphere samples. PERMANOVA confirmed statistically significant differences between compartments, and ANCOM-BC2 identified differentially abundant bacterial families across compartments. Together, these results indicate that compartment identity is a major factor associated with bacterial community composition in *C. colocynthis*.

Soil represented the most diverse compartment and showed a distinct community composition compared to plant-associated compartments. This pattern is consistent with the role of soil as a broad microbial reservoir (59). The higher representation of taxa such as Chloroflexota and other soil-associated groups further supports this distinction. These observations align with the general concept that rhizospheric soil provides a diverse microbial pool from which plant-associated compartments recruit a subset of taxa (60). In semi-arid environments, soil physicochemical constraints, water limitation, and nutrient availability may further influence microbial community assembly processes (21, 61). Multivariate analyses (CAP/db-RDA) indicated that measured environmental variables were associated with community variation, particularly in the second-year sampling group. However, relatively low to moderate explained variance suggests that additional factors, including host-related selection and unmeasured environmental heterogeneity, may also contribute to community structure.

Root-associated communities showed intermediate diversity and composition between soil and leaf endosphere compartments. Several bacterial families enriched in roots, including Microbacteriaceae, Rhizobiaceae, Devosiaceae, Pseudomonadaceae, and Xanthomonadaceae, are commonly reported in plant-associated environments. These taxa include members previously associated with plant colonization, nutrient cycling, and stress tolerance (62). In particular, Pseudomonadaceae, including *Pseudomonas*, contains strains described for siderophore production, hormone modulation, phosphate solubilization, ACC deaminase activity, and biocontrol potential (63, 64). Actinomycetota-related families, such as Microbacteriaceae, Micrococcaceae, Nocardioidaceae, and Micromonosporaceae, are frequently associated with soil–root interfaces and have been linked to secondary metabolite production and adaptation to dry environments (65).

Leaf endosphere communities showed lower diversity compared to soil and root compartments and formed distinct clusters in beta-diversity analyses. This pattern is consistent with stronger environmental filtering in the leaf endosphere, where microbial communities are exposed to limited nutrients, desiccation, UV radiation, and host-driven selection (66, 67). The recurrent detection of Pseudomonadaceae and *Pseudomonas* across plant-associated compartments suggests that these taxa may include members adapted to both root and aerial plant habitats, although their ecological roles likely vary at the strain level. *Pseudomonas* is widely recognized as a key plant-associated genus with strong capacities for nutrient mobilization and plant growth promotion, including siderophore production, phosphate solubilization, phytohormone-mediated growth regulation, and biocontrol activities that contribute to plant health and ecological resilience in host–microbe systems (63).

The observed compartmental structure is generally consistent with previous microbiome studies on *C. colocynthis* in arid and semi-arid environments. Previsou investigation has reported diverse bacterial assemblages across soil and plant compartments, including taxa commonly associated with soil functioning and plant-associated lifestyles such as Acidobacteria, Bacteroidetes, and Actinobacteria (29). In the present study, the rhizosphere soil again represented the most diverse compartment, while root and leaf endosphere communities appeared more strongly shaped by filtering processes. Differences in dominant taxa compared to previous studies are expected due to variation in geography, soil properties, climate conditions, host genotypes, and methodological choices such as 16S rRNA gene region selection (68). These differences highlight the context dependency of microbiome assembly in stress-adapted plant species.

The culture-dependent approach provided complementary resolution to amplicon-based profiling by enabling the recovery of cultivable bacterial taxa (69, 70). Sanger-based identification indicated that the cultured collection included several dominant genera, with *Achromobacter*, *Pseudomonas*, and *Glutamicibacter* among the most frequently detected. Phylogenetic analysis of 16S rRNA gene sequences confirmed the taxonomic diversity of the isolates and provided isolate-level resolution complementary to the amplicon dataset. Most cultured genera were also detected in the sequencing data, indicating overlap in taxonomic detection between approaches, although not necessarily at the strain level.

The cultured isolate collection provides an important bridge between the amplicon-detected community and cultivable members, consistent with the complementary use of culture-dependent and culture-independent approaches in microbiome studies (69–71). Sanger-based identification showed that the cultured collection included several genus-level groups, with *Achromobacter*, *Pseudomonas*, and *Glutamicibacter* being the most represented. The representative 16S rRNA gene phylogeny confirmed the taxonomic breadth of the collection, while the complete tree provided isolate-level context for the GenBank-submitted dataset. Importantly, most cultured genera were also detected in the amplicon dataset. This overlap reflects taxonomic correspondence rather than detection of the same strain by both approaches.

Previous studies have reported that *C. colocynthis* can host bacterial isolates with plant-associated functional traits, including antimicrobial activity, pathogen suppression, and plant growth promotion. Endophytic actinobacteria such as Streptomycetaceae and Nocardiopsaceae have been associated with antibacterial activity (28). Other isolates have shown inhibitory effects against phytopathogens such as *Fusarium solani* and *Pythium aphanidermatum*, as well as growth-promoting effects in model plants (72). Additional studies have reported traits such as phytohormone production and mineral solubilization in rhizospheric isolates from arid environments (72). These findings suggest that cultivable members of the *C. colocynthis* microbiome may include bacteria with functional potential, although these traits remain strain-dependent and require experimental validation.

Genome-informed functional profiling of selected isolates provided additional insight into potential ecological roles. Whole-genome sequencing revealed the presence of genes associated with stress response and environmental adaptation, consistent with genome-based approaches for predicting plant-associated traits (73). Genes related to oxidative stress response were widely distributed among isolates, which is consistent with exposure to semi-arid environmental conditions and with the role of plant-associated bacteria in abiotic stress tolerance (74, 75). Osmoprotection-related genes were also frequently detected, suggesting potential relevance to water limitation and osmotic stress adaptation.

Functional genes associated with siderophore production, nitrogen metabolism, ACC deaminase activity, motility, chemotaxis, biofilm formation, secretion systems, and antagonistic interactions were detected with variable distribution across isolates. These functions are commonly associated with plant-associated bacteria, although their expression is context-dependent and may vary with environmental conditions and host interactions (63, 76). The observed variability suggests functional heterogeneity among cultivable isolates rather than a uniform functional profile.

Overall, the combined dataset indicates a structured bacterial community associated with *C. colocynthis*, differentiated across soil, root, and leaf endosphere compartments. Soil acts as a diverse microbial reservoir, roots represent a selective transition zone, and the leaf endosphere appears as a more strongly filtered habitat. The partial overlap between cultured isolates and amplicon-detected taxa, together with genome-encoded functional traits related to stress adaptation and plant association, suggests that the cultivable fraction includes bacterial lineages with potential ecological roles in semi-arid environments.

## Conclusion

This study demonstrates that *C. colocynthis* hosts a structured and compartmentalized bacterial microbiome primarily shaped by habitat differentiation across the rhizosphere, roots, and the leaf endosphere. Rhizosphere functions as a highly diverse microbial reservoir, while root-associated and leaf endosphere communities appear progressively more selective, consistent with increasing environmental and host-mediated filtering.

By integrating amplicon sequencing, culture-dependent isolation, and genome-informed functional profiling, the study reveals a general concordance between taxonomically detected and cultivable bacterial groups. It also highlights the prevalence of ecologically relevant taxa, including *Pseudomonas*, *Achromobacter*, and several Actinobacteria-related lineages. Functional genomic analyses further indicate that many isolates encode traits associated with stress tolerance, nutrient acquisition, and plant interaction, which is consistent with adaptation to semi-arid environmental conditions. These findings suggest that the *C. colocynthis* microbiome constitutes a structured and functionally diverse bacterial network potentially contributing to plant persistence in harsh environments. The cultivable fraction represents a particularly promising reservoir for identifying bacteria with ecological and biotechnological relevance in arid and semi-arid systems. Future work should focus on experimentally validating the functional potential of key isolates under environmentally relevant conditions, including plant inoculation assays to assess growth promotion and stress mitigation. In addition, the development of synthetic bacterial consortia based on functionally complementary strains could provide a more realistic framework for improving plant performance under drought and nutrient-limited conditions. Metabolomic and transcriptomic profiling of both plants and associated bacteria would further help to clarify microbe–plant interaction mechanisms and identify key bioactive compounds involved in stress adaptation. Integrating these approaches with field-based trials across environmental gradients would also strengthen the ecological relevance and translational potential of the findings for sustainable agriculture in semi-arid ecosystems.

## Acknowledgments

We thank Prof. Khaoula Errafii, Dr. Safaa Machraoui, and Mr. Abdelhadi Ziami for their valuable technical assistance and support. We gratefully acknowledge the financial support provided by OCP Nutricrops through Project AS-85. We also acknowledge the Toubkal Supercomputer Team at UM6P (Morocco) for providing computational resources and technical support that contributed to this work.

## Author Contributions

K.A.S.M. contributed to sampling, experimental work, sequencing data analysis, and manuscript drafting. S.M. and I.K. assisted with laboratory work and sample preparation. N.R. contributed to sequencing and data curation. F.A. contributed to library preparation. H.M. contributed to conceptualization, manuscript drafting, review, and supervision. All authors contributed to the manuscript and approved the submitted version.

## Data Availability

The 16S rRA gene amplicon sequencing datasets generated in this study are available in the NCBI Sequence Read Archive under accession numbers listed in Table S11. Sanger sequences were deposited at the NCBI and accession numbers are listed in Table S10.

